# Whole-genome SNP analysis elucidates the genetic population structure and diversity of Acrocomia species

**DOI:** 10.1101/2020.10.08.331140

**Authors:** Brenda G. Díaz, Maria I. Zucchi, Alessandro. Alves-Pereira, Caléo P. de Almeida, Aline C. L. Moraes, Suelen A. Vianna, Joaquim A. Azevedo-Filho, Carlos A Colombo

**Author notes:** Corresponding author (CAC).

## Abstract

Acrocomia (Arecaceae) is a genus widely distributed in tropical and subtropical America that has been achieving economic interest due to the great potential of oil production of some of its species. In particular *A. aculeata*, due to its vocation to supply oil with the same productive capacity as the oil palm even in areas with water deficit. Although eight species are recognized in the genus, the taxonomic classification based on morphology and geographic distribution is still controversial. Knowledge about the genetic diversity and population structure of the species is limited, which has limited the understanding of the genetic relationships and the orientation of management, conservation, and genetic improvement activities of species of the genus. In the present study, we analyzed the genomic diversity and population structure of seven species of Acrocomia including 117 samples of *A. aculeata* covering a wide geographical area of occurrence, using single nucleotide Polymorphism (SNP) markers originated from Genotyping By Sequencing (GBS). The genetic structure of the Acrocomia species were partially congruent with the current taxonomic classification based on morphological characters, recovering the separation of the species *A. aculeata, A. totai, A. crispa* and *A. intumescens* as distinct taxonomic groups. However, the species A. media was attributed to the cluster of *A. aculeata* while *A. hassleri* and *A. glauscescens* were grouped together with *A. totai*. The species that showed the highest and lowest genetic diversity were A. totai and A. media, respectively. When analyzed separately, the species A. *aculeata* showed a strong genetic structure, forming two genetic groups, the first represented mainly by genotypes from Brazil and the second by accessions from Central and North American countries. Greater genetic diversity was found in Brazil when compared to the other countries. Our results on the genetic diversity of the genus are unprecedented, as is also establishes new insights on the genomic relationships between Acrocomia species. It is also the first study to provide a more global view of the genomic diversity of *A. aculeata*. We also highlight the applicability of genomic data as a reference for future studies on genetic diversity, taxonomy, evolution and phylogeny of the Acrocomia genus, as well as to support strategies for the conservation, exploration and breeding of Acrocomia species and in particular *A. aculeata*.

## Introduction

The genus Acrocomia is endemic to tropical and subtropical America. This genus is one of the most taxonomically complex concerning species in the family Arecaceae [1]. Taxonomic classifications of Acrocomia are mostly limited to the description of species based on morphological and geographical distribution information. However, extensive morphological plasticity, especially for species with wide geographical distribution, has hindered the taxonomic resolution of species. Since the description of the genus Acrocomia by Martius in 1824 [2], many species have been included and removed from the genus. From the most recent classifications, Henderson et al. [3] attributed only two species to the genus. One is *A. aculeata* (Jacq.) Lodd. Ex Mart., which is large (arboreal) and widely distributed throughout Central, North, and South America. The other is *A. hassleri* (Barb. Rodr.) WJ Hahn, which is small in size and is restricted to the Cerrado savanna in Brazil and part of Paraguay. Lorenzi et al. [4] recognized seven species for the genus. Six of these are found in Brazil: *A. aculeata, A. intumescens*, and *A. totai* have an arboreal size and are mainly differentiated by the stipe characteristics. *A. hassleri, A. glaucescens*, and *A. emensis* are small size and are differentiated by their height. The seventh species, *A. crispa*, has an arboreal size and is endemic to Cuba. The Plant List [5] and The Palmweb [6] recognized *A. media* as the eighth species. It is endemic to Puerto Rico. Therefore, the systematics of the genus Acrocomia remain controversial, with the number of species not well resolved and very few studies having addressed species delimitation, population genetic diversity and structure, and inter-species relationships.

*A. aculeata, A. totai*, and *A. intumescens* are the species of greatest economic interest, mainly due to their many applications and products obtained, with practically all parts of the palms used. The fruits are important for the production of vegetable oil as a bioenergy source and flour for human and animal consumption [7] as well as for medicinal uses [7, 8]. Of these three species, *A. aculeata* is distinguished by its high productive capacity and oil quality [9]. The oil production of 4,000 oil L/ha/year estimated in Brazil far surpasses soybeans (400 L/ha) [9] and equals the oil palm, which is considered the oilseed with the highest oil yield per area, with an oil production volume of up to 6,000 L/ha [10, 11].

*A. aculeata* is an arborescent heliophile and monoecious. This species produces unisexual flowers in the same inflorescence [3, 4]. It has a mixed reproductive system, with a preference for allogamy [12]. It is a diploid species (2n = 30), with a genome size of 2.8 Gbp [13]. *A. aculeata* has a wide geographic distribution, occurring naturally from northern Mexico and the Antilles to southern Brazil [3, 4, 14, 15]. It is commonly found in savanna areas, but also is found in tropical and subtropical forests, and in the dry forests of Caatinga [3, 4, 16] and has adapted to sandy soils and regions with low water availability [17]. Besides being a perennial species, it is beneficial for soil management and conservation since its useful life can exceed 50 years. Colombo et al. [9] identified *A. aculeata* as a promising resource for sustainable large-scale production of vegetable oil.

Although the economic interest in some Acrocomia species is growing, little is known about infrageneric relationships, levels of genetic diversity and structure, and patterns of gene flow at the genus level. The population genetics approach can assist in species delimitation and provide reference information on the genetic diversity and structure within and between species. Such knowledge is essential for more efficient management and economic exploration of the species and can guide strategies for domestication and conservation of these genetic resources. *A. aculeata* is an emerging crop with incipient domestication. The analysis of genetic diversity of *A. aculeata* is crucial to guide the selection of the most promising materials for crop use, to maximize genetic gains, and to more effectively contribute to the creation of commercial cultivars.

In this context, molecular markers have been broadly adopted in plants as an essential tool to investigate genetic diversity in ecological, phylogenetic, and evolutionary studies. In addition, they have been widely used for direct management, conservation, and genetic breeding of several species [18]. More recently, next-generation sequencing (NGS) has facilitated the identification of single nucleotide polymorphisms (SNPs), which have emerged as the most extensively used genotyping markers due to their abundance and distribution in the genome. The use of SNPs has considerably expanded knowledge of the genetic diversity of genomes of various plant species [19] at low cost and without the need for reference genomes [20–22]. However, SNPs have not been used as markers in genetic studies of Acrocomia species.

In *Acrocomia*, microsatellites or simple sequence repeats (SSR) have been the most used molecular markers, with the main objective of evaluating the genetic diversity and structure of natural populations and germplasm banks [12, 23–26]. Other approaches include the use of internal transcribed ribosomal 18S-26S spacer (ITS region) [27] and random amplification of polymorphic DNA (RAPD) markers [28]. However, most studies have focused on *A. aculeata* [12, 23, 25, 26, 29]. Only one study has analyzed the genetic diversity of *A. totai* (Lima et al., 2020).

Considering the wide distribution of *A. aculeata* in the Americas, all the studies carried out using molecular markers have revealed a limited panorama of species genetic diversity because they considered a very small geographic sampling, with genotypes obtained mainly from the states of São Paulo and Minas Gerais in Brazil (Abreu et al., 2012; Lanes et al., 2015; Mengistu 2016; Oliveira et al., 2012; Coelho et al., 2018). Only a single study has evaluated the genetic diversity of natural populations of *A. aculeata* (termed *A. mexicana)* from another country besides Brazil, that being Mexico [30].

Faced with the difficulties of taxonomic resolution of the genus Acrocomia, our study aimed to apply a population genomic approach to elucidate the genetic diversity and genetic relationships of species through genomic polymorphism data. Recognizing the increasing importance of *A. aculeata* and the lack of genetic reference data in the different countries where it grows, we analyzed the genetic diversity and structure of natural populations of *A. aculeata*, considering its wide occurrence in the American continent.

The present study is unprecedented because it was conducted using seven Acrocomia species and a wide sampling of *A. aculeata* from several countries in the American continent. This is the first study carried out with SNP markers for the genus.

## Material and Methods

### Plant material and DNA extraction

In the present study, we considered 172 samples to represent seven from eight Acrocomia species: *A. aculeata, A totai, A. intumescens, A. media, A. crispa, A. hassleri, A. glaucescens*. The samples were obtained from different locations in order to represent the entire geographic distribution described in the literature for the respective species[3, 4]. The species *A. aculeata*, with a greater distribution in America, was represented by samples from five countries (Fig 1 and S1 Table).

**Fig 1.**
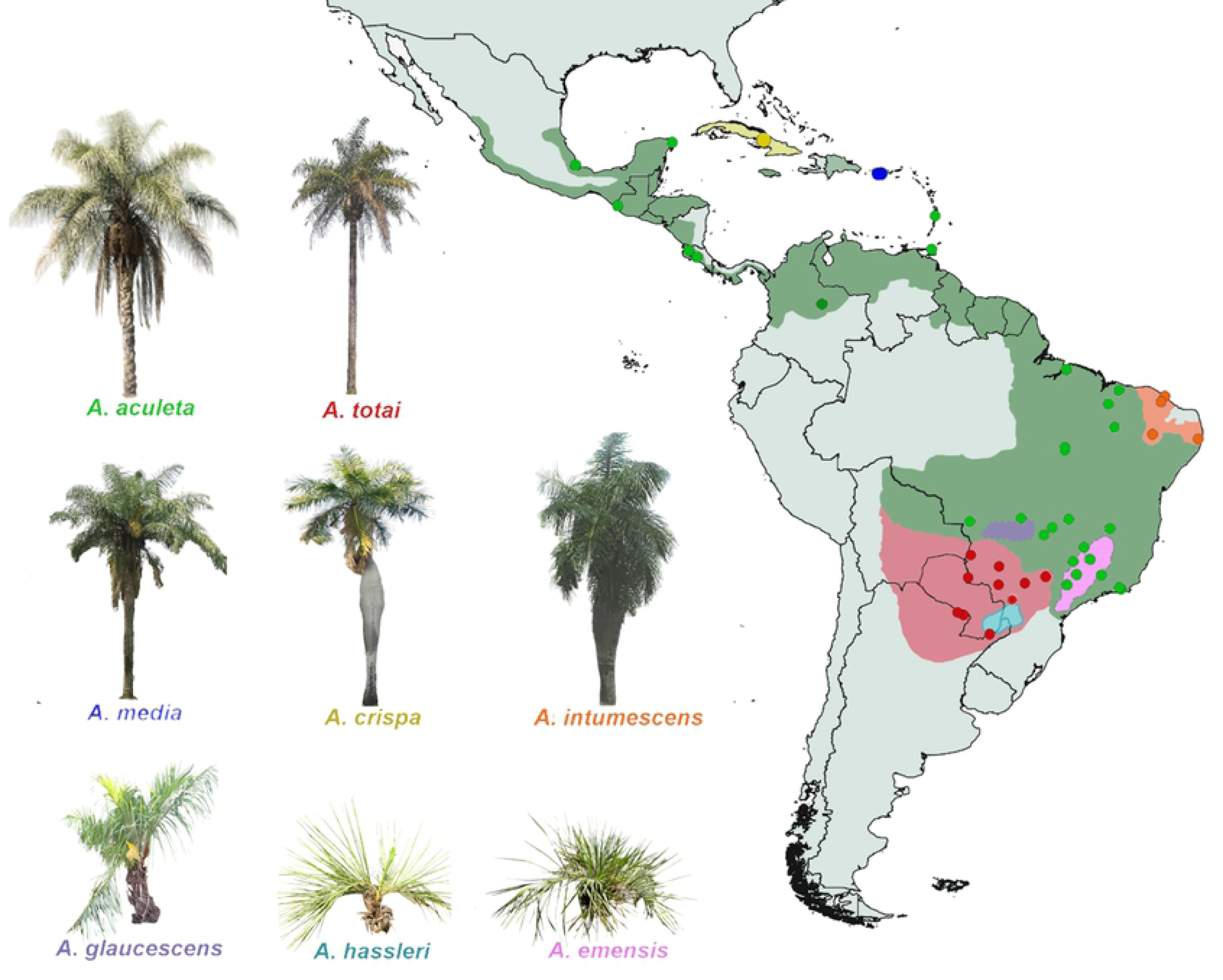
Schematic map of Acrocomia species distribution and geographic location and origin of samples. Data used to generate the species distribution (Colored shading) are based on occurrence record data from GBIF (Global Biodiversity Information Facility www.gbif.org) and Lorenzi et al., [4]. Circles represent geographical location and origin of samples in this study. Image sources: *A. aculeata, A. totai, A. hassleri, A. glaucescens. A. emensis* (B. G. Díaz); *A. intumescens*; *A. media* and *A. crispa* (S. A. Vianna.)

The total genomic DNA was extracted from leaf material using the Doyle & Doyle [31] protocol. We evaluated the quality and quantity of DNA on a 1% agarose gel, on the NanoVue ™ Plus spectrophotometer (GE Healthcare), and through fluorescence using the QubitTM dsDNA BR Assay (Qubit - Life Technologies). Based on the obtained reading, we standardized the DNA to a concentration of 30ng.μl-1.

### GBS library preparation and High-Throughput Sequencing

To obtain SNPs, we developed genomic libraries using the GBS (Genotyping by Sequencing) technique according to the protocol described by Poland et al. [32], with modifications. We digested 7 µl of the genomic DNA [30ng.μl-1] from each sample at 37 ° C for 12 hrs with the enzymes *NsiI* and *MspI*. Subsequently, 0.02 µM of specific adapters for the Illumina technology (containing the barcode sequences and complementary to the Illumina ™ primers for sequencing) were connected to the fragments ends generated in the digestion. The ligation reaction was carried out at 22°C for 2 h; 65°C for 20 min; 10°C indefinitely.

After adapters ligation, we purified the samples using QIAquick PCR Purification Kit (Qiagen). The library was enriched by PCR. We performed eight replicates, each one containing 10 µL of purified and amplified ligation, using 12.5 µL of Phusion® High-Fidelity PCR Master Mix NEB (New England Biolabs Inc.), and 2 µl of Illumina forward and reverse [10 µM] primers ™, in a final volume of 25 µL, using the following amplification program: 95°C for 30 s, followed by 16 cycles of 95°C for 10 s, 62°C for 20 s, 72°C for 30 s, ending at 72°C for 5 min. Finally, we purified the library using QIAgen’s QIAquick PCR Purification Kit.

The verification of average size of the DNA fragments using the Agilent DNA 12,000 kit and the 2100 Bioanalyzer System (Agilent) equipment. The libraries were quantified by qPCR using the CFX 384 real-time thermocycler (BioRad) with the aid of the KAPA Library Quantification kit (KAPA Biosystems). We prepared two libraries of 96 samples each, which were sequenced using Illumina’s NextSeq 500/550 Mid Output Kit v2.5 (150-cycle), on the NextSeq550 platform (Illumina Inc., San Diego, CA).

### SNP identification

We performed the identification of SNP markers using the Stacks *v*. 1.42 *pipeline* [33]. We used the *process_radtags* module to demultiplex the samples and to remove the low-quality readings. As there is no reference genome for Acrocomia, we aligned the sequences and organized the loci using the *ustacks* module with the following parameters: the minimum sequencing depth (-m) ≥ 3, the maximum distance between *stacks* (-M) = 2; and the maximum distance between primary and secondary sequences (-N) = 2. Subsequently, a locus catalog was built using the *cstacks* module, allowing a maximum of 2 differences between stacks (-n) from different individuals. We eliminated loci with lower values of probability (lnl_lim -10) by the *rxstacks* correction module. The SNPs were filtered using the *populations* module, retaining only one SNP per sequence, with a minimum depth of 3X sequencing, minor allele frequency ≥ 0.01, and minimum occurrence in 75% of individuals in each location/population. After filtering, we identified 3269 SNPs (S1 File).

### Identification of putatively neutral and under selection loci

We identified neutral SNPs and loci putatively under selection (outliers). To reduce the possibility of identifying false positives, we applied three approaches to identify outlier loci. For the first approach, we used the method based on Principal Component Analysis (PCA) from the *pcadapt* package [34], on the R platform [35]. The *pcadapt* method assumes that SNPs excessively related to the population structure are candidates to be under adaptive selection. In this approach, no a priori information about the number of populations was introduced. Initially, we carried out the principal component analysis (PCA) to define the structure of the data set, adopting the Mahalanobis distance from the z-scores in the first k-components of each locus to identify the most related loci to the population structure. In the second approach, we used the *fsthet* package [36] based on Wright’s F_ST_ fixation index [37] to identify the loci with deviation from the expected relationship between F_ST_ and heterozygosity (H_E_), using the island migration model [38].

The third approach we adopted to test the association of environmental variables with the genetic variation of SNP markers was the LFMM (Latent Factor Mixed Models) [39], using the LEA package (Landscape Genomics and Ecological Association Test) [40] on R. platform [35]. We used nineteen bioclimatic variables related to precipitation and temperature, in addition to the minimum, average and maximum values of wind speed, vapor pressure, and solar radiation, obtained from the WorldClim database [41]. We performed the analyzes with the following variables (correlation ≤0.8): average annual temperature, average daytime variation, isothermality, average temperature of the wettest four-month period, annual temperature variation, annual precipitation, precipitation in the driest month, precipitation seasonality, radiation maximum solar radiation, minimum solar radiation, and average wind speed. For the lfmm function, five replicates were performed with 200,000 MCMC interactions after 50,000 burn-ins. For the association tests, the genetic structure presented between the individuals was considered with the SNMF analysis [39], determining the most likely number of genetic groups for the different data sets, using 100,000 MCMC interactions, and 10 repetitions□ for the number of groups (K) varying between 1 to 15. The LFMM analysis considered K = 8 (species) and 6 (*A. aculeata* Americas). The associations with environmental variables were identified for the loci with corrected p-values, considering FDR = 0.1 of environmental variable association to detect SNPs, from the LEA package [40], using the sparse non-negative matrix factorization function (snmf) [42]. We carried out the identification of environmental variables by the principal component analysis, adopting 19 bioclimatic variables from the WorldClim database [41], and selecting the variables that showed the highest correlation. For the snmf function, the most likely number of populations for the different data sets was determined using 100,000 interactions, and 10 repetitions □ for K = 1-15.

The identification of SNPs hypothetically under selection (outliers) was performed for the following groups: 1) In the genus Acrocomia, considering the species as groups, and 2) within *A. aculeata*, considering as groups the samples’ countries of origin. We considered as loci putatively under selection those shared between the three identification methods (*fsthe*t, *pcadapt* and LFMM) (S2 Table). Consequently, we adopted the remaining SNPs considered neutral for the analysis of population genomic diversity and structure.

### Population structure

We used all samples (S1 Table) to perform the analysis of the genomic structure for de Acrocomia genus and to infer the number of the most likely groups using the software Structure v.2.3.4 [43], considering only neutral SNPs (3227). We also used the same software to access the genomic structure of *A. aculeata* separately, considering 3259 neutral SNPs identified for the species. Each analysis in Structure was performed with a burn-in of 100,000 interactions, followed by 500,000 repetitions of the Markov Chain Monte Carlo (MCMC) in 10 independent simulations, and without prior information to define the clusters. The number of clusters (K) was determined using the average likelihood values of the ΔK method [44] implemented in the program Structure Harvester [45]. The participation coefficient for each access was given by the alignment of five repetitions of the best K through the CLUMPP method [46] by the software CLUMPAK [47].

To visualize the genetic relationships among Acrocomia species and within the *A. acuelata*, we obtained the Nei genetic distance [48] between the individuals of each data set, and the Neighbor-Joining (NJ) hierarchical classification method with 20000 bootstrap repetitions, using the poppr package[49] on R [35].

In addition, the Principal Component Analysis (PCoA) was also carried out through the ADE 4 package [50]to explore the genetic structure of the different groups using only neutral SNPs, and was visualized graphically by the ggplot2 package [51].

### Analysis of genomic diversity

We conducted the population diversity analysis only with the SNP data set identified as neutral for two groups or taxonomic levels: 1) The genus Acrocomia (except the species *A. hassleri* and *A. glaucescens* as they contain only one individual for each species), and 2) *A. aculeata*. Population estimates of allelic richness, percentage of total alleles by locus, observed heterozygosity, expected heterozygosity, and inbreeding coefficient were calculated using the diveRsity [52], poppr [49], and the PopGenKit packages [53] on R platform [35]. To minimize the effect of differences in the number of samples of each population, we calculated the allelic richness (Ar) and the richness of private alleles (ap) for populations of each group or taxonomic level, by the rarefaction method implemented in the software HP-Rare v.1.1 [54].

## Results

In population genetics, neutral loci are genomic regions that are influenced by mutational dynamics and demographic effects, and not by selection. However, loci under selection (i.e., outliers) generally behave differently and therefore reveal “extreme” patterns of variation [55, 56]. Since most population genetic inferences are based on neutral loci, the loci under selection can greatly influence the estimates of genetic parameters. In this sense, it is important to identify and remove the outlier loci from the analysis, with the aim to infer more reliable parameters of population genetic diversity and structure.

Based on pcadapt, fsthet, and LEA, we identified 42 outlier loci for all samples or taxonomic groups for the genus Acrocomia, and 10 outlier loci for the taxonomic group formed by samples of *A. aculeata*. The neutral datasets for the different groups were constructed by removing the outliers. After the removal of outlier loci (S2 Table), genus Acrocomia (all species) and *A. aculeata* contained 3227 and 3259 neutral loci, respectively.

### Genomic structure of Acrocomia spp

Structure 2.3.4 software [43] was initially used to access the genomic structures of 172 samples of Acrocomia species based on 3227 neutral SNPs. ΔK had a maximum value of K = 7 (S1 Fig), indicating the existence of seven genetic groups (Fig 2). Samples with an attribution probability score > 0.75 and < 0.75 were assigned to the “pure group” and “admixture group”, respectively. Based on the classification of Lorenzi [4] and the geographic distribution of the species, we observed a substructure of samples considered to be *A. aculeata*. Two well-defined subgroups (clusters 1 and 3) strongly associated with the geographical origin of the samples were evident. Cluster 1 (Fig 2) was composed of 38 samples of *A. aculeata* from Central and North America (Costa Rica, Trinidad and Tobago, Puerto Rico, and Mexico) and Colombia. Cluster 2 (Fig 2) comprised 39 samples of *A. totai* and five samples considered as *A. aculeata*. Of the latter, four were collected in the state of Parana, southeastern Brazil, (XAM, PR) and one in state of Tocantins, northern Brazil (PAL). The samples from Campo Grande (CGR) showed low mixture levels with clusters 1 and 5 of *A. aculeata*. Cluster 3 (Fig 2) consisted of 39 samples from Brazil. The majority (n = 34) of these samples were from the southeast region of the country, with five from the north region (BEL population). Cluster 4 was exclusively *A. crispa* samples, with a 100% probability of assignment to the cluster.

**Fig 2.**
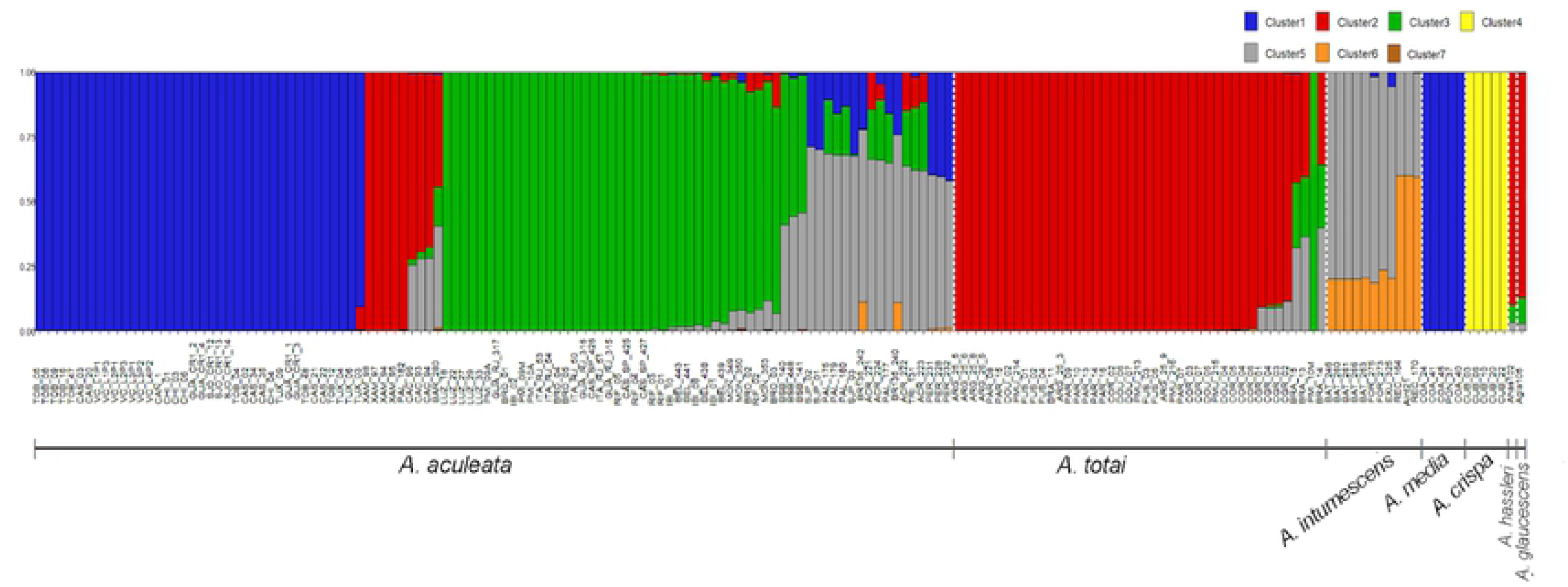
Genomic structure of 172 samples from Acrocomia species based on 3227 neutral SNPs loci. The y-axis is the population membership, and the x-axis is the sample. Each vertical bar represents a sample and color represent separate clusters (k =7)

Based on assignment probabilities ≤ 0.75, some samples were assigned to an admixture group. Twenty samples of *A. aculeata* from the central-west, north, and northeast regions of Brazil, and all samples of *A. intumescens* displayed a similar genomic composition, with a median level of assignment (≥ 0.50). A genetic admixture of *A. aculeata* samples in cluster 5 (Fig 2) with samples mainly from clusters 1 and 3 was evident. *A. intumescens* samples presented a mixture of clusters 5 and 6, with cluster 6 being practically exclusive to the species. Individuals from Cáceres, MT (CAC), and Braúna, São Paulo (SP) (BRA), with a greater assignment to cluster 2, also showed a significant degree of admixture with clusters 3 and 5.

The NJ and PCoA analyses (Figs 3a and 3b) performed with all the samples showed strong agreement with the results of the Bayesian analysis performed using Structure software. However, the NJ tree showed higher resolution in group/cluster recovery than the PCoA. In both analyses, *A. crispa* was clearly separated from the rest of the Acrocomia species. In addition, there is a clear genomic differentiation between *A. aculeata* and *A. totai*. Similar to the results obtained using the Structure software, the NJ analysis also recovered the substructure within *A. aculeata*, separating the Brazilian samples from those from other countries (Fig 3a). This separation did not result from the PCoA (Fig 3b). In agreement with the results obtained using the Structure software, both PCoA and NJ grouped *A. media* and *A. intumescens* samples into the cluster formed mainly by *A. aculeata*, with *A. hassleri* and *A. glaucescens* grouped into the *A. totai* cluster. The results of NJ and PCoA also agreed concerning the allocation of samples from Xambré, PR (XAM) originally considered as *A. aculeata* in the cluster of *A. totai*. Samples from Braúna, SP (BRA) and Cáceres, MT (CAC), which were identified as an admixture by the Structure software, occupied an intermediate position between the clusters formed mainly by *A. aculeata* and *A. totai* in the PCoA.

**Fig 3.**
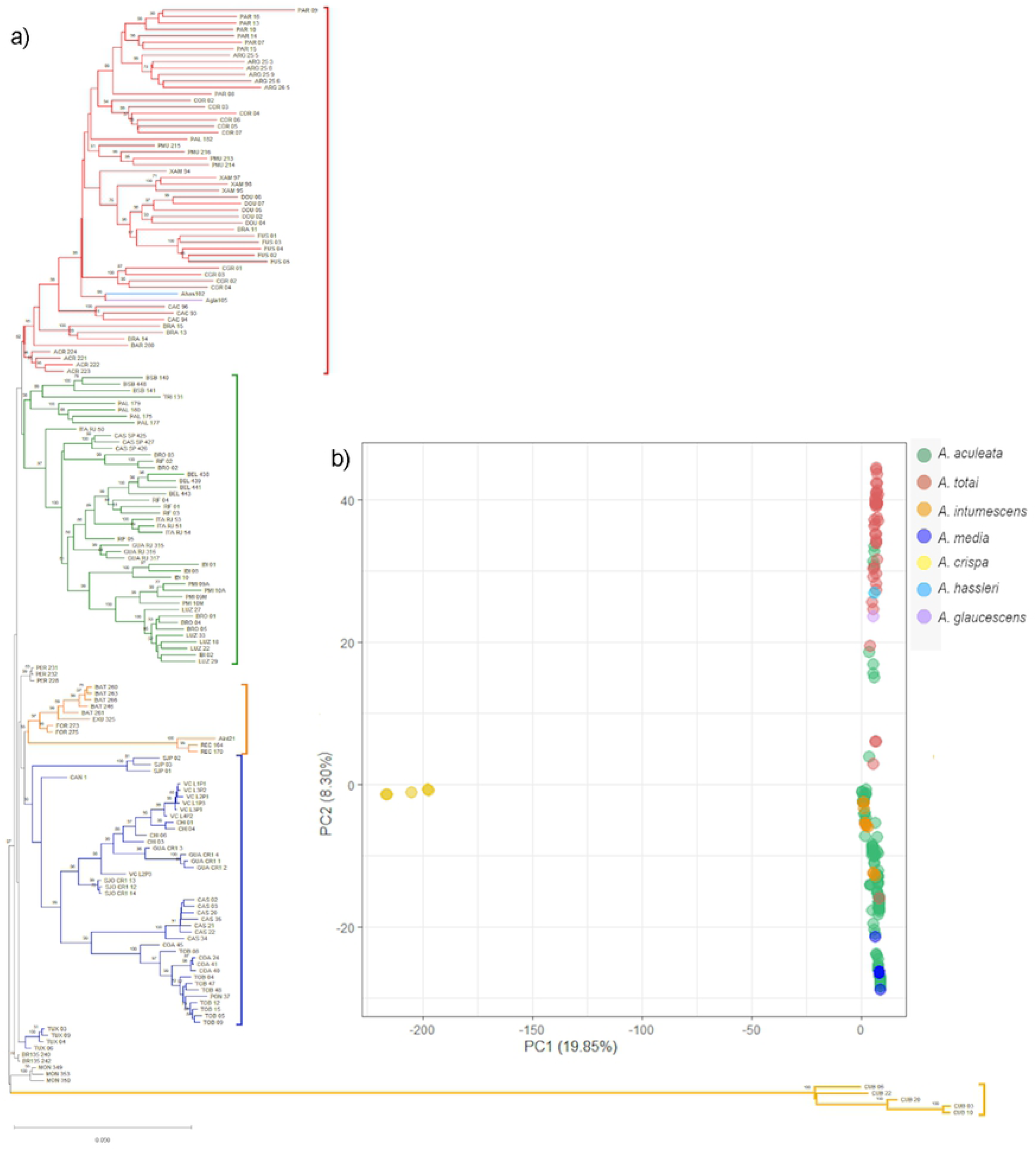
Neighbor-joining (NJ) tree and principal components analysis (PCoA) of Acrocomia species. a) Scatterplot of the principal components analysis (PCoA) showing the dispersion of samples across the first two principal components and b) Neighbor-Joining dendrogram based on Nei’s genetic distance. Bootstrap support of nodes is shown.

Based on the Structure software results (Fig 2) and NJ and PCoA data (Fig 3a and 3b), the samples from Xambrê, PR (XAM) previously considered *A. aculeata* were treated as *A. totai* species for further analysis of differentiation and genomic diversity. The F_ST_ values enabled a moderate genetic differentiation between species, with an average value of 0.469. The F_ST_ values between species (Table 1) ranged from 0.083 (*A. aculeata* vs. *A. totai)* to 0.946 (*A. media* vs. *A. crispa*). In agreement with the genomic structure analysis findings, all comparisons between *A. crispa* and the other species showed higher values of F_ST_, demonstrating a greater degree of genetic differentiation of *A. crispa* with the other species.

**Table 1.**
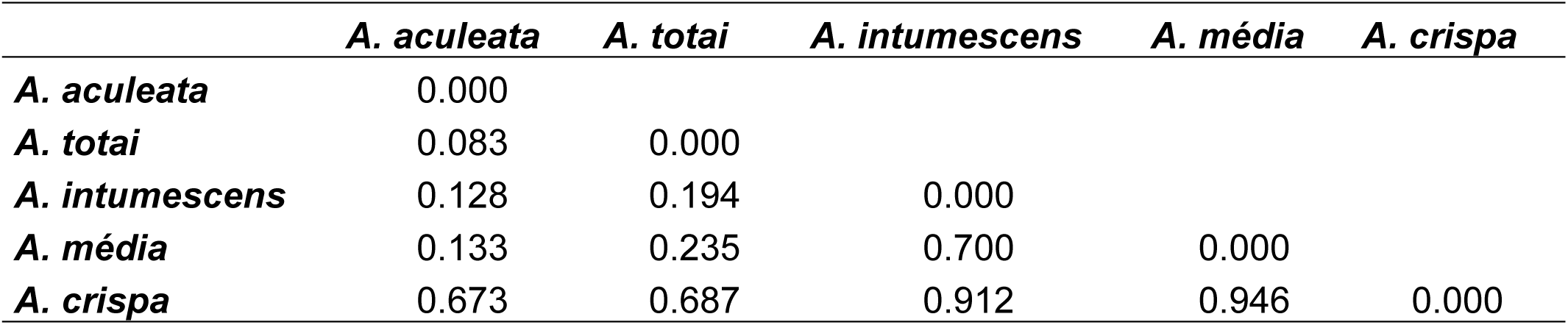
Pairwise FST estimates among five species of Acrocomia

### Genomic diversity between species

The number of polymorphic loci of the five Acrocomia species ranged from 0.017 to 0.601. *A. aculeata* had the highest mean and *A. media* had the lowest mean (Table 2). The genomic diversity based on the average expected heterozygosity (H_E_) in the species ranged from 0.106 in *A. totai* to 0.005 in *A. media*. However, *A. crispa* was the species with the highest allelic richness (2.29) and the highest allelic richness of private alleles (0.17), while *A. media* presented the lowest values of allelic richness and allelic richness of private alleles (1.08 and 0.01, respectively). The inbreeding coefficient (F) values were high for all species, indicating relatively high levels of inbreeding in Acrocomia species, with the exception of *A. media*, which presented negative values (Table 2).

**Table 2.** Genetic diversity parameter estimates for Acrocomia species calculated from 3227 neutral loci SNPs.

| Species | Na | Ne | I | H<sub>O</sub> | H<sub>E</sub> | Ar | PAr | f |
| --- | --- | --- | --- | --- | --- | --- | --- | --- |
| *A. aculeata* | 1.601 | 1.142 | 0.157 | 0.031 | 0.093 | 1.20 | 0.03 | 0.479 |
| *A. totai* | 1.534 | 1.160 | 0.176 | 0.074 | 0.106 | 1.23 | 0.07 | 0.262 |
| *A. intumescens* | 1.053 | 1.027 | 0.037 | 0.011 | 0.025 | 1.10 | 0.02 | 0.483 |
| *A. média* | 0.993 | 0.984 | 0.007 | 0.006 | 0.005 | 1.08 | 0.01 | -0.145 |
| *A. crispa* | 0.630 | 0.619 | 0.028 | 0.006 | 0.020 | 2.29 | 0.21 | 0.591 |

Mean of different alleles (Na), effective allele (Ne), Shannon's Index (I), Observed (H<sub>O</sub>) and Expected (H<sub>E</sub>) Heterozygosity, allelic richness (Ar), private alleles richness (PAr) and Fixation index (f)

### Genomic structure of *A. aculeata*

The population structure of all the *A. aculeata* samples was evaluated using 3259 hypothetically neutral SNPs. Using the method of Evanno [44] the most probable Δk was K = 2 (S2 Fig). This finding supported the presence of two genetically distinct subpopulations previously identified in the structure analysis at the genus level (Fig 2). The two groups were mainly associated with geographical origin, given that samples from Central and North America (Colombia, Costa Rica, Trinidad and Tobago, and Mexico) were grouped in cluster 1, and most of the collected in Brazil were grouped in cluster 2 (Fig 4).

**Fig 4.**
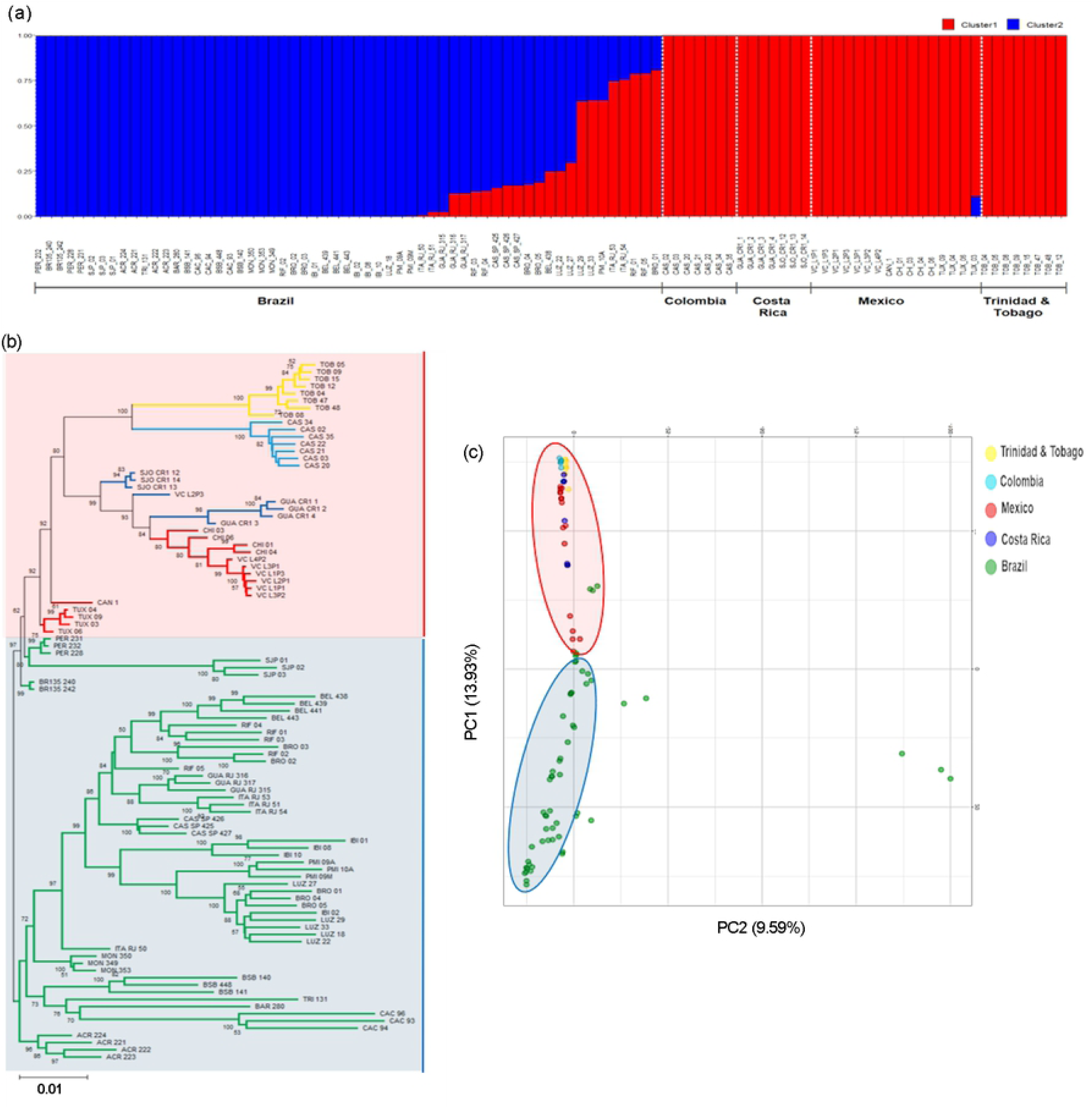
Population genomic structure within *A*. *aculeata*, based on 3256 neutral loci SNPs. a) Genomic structure from Bayesian analyses (k =2). The y-axis is the population membership, and the x-axis is the sample. Each bar represents an individual and each color is inferred membership in each of the cluster; b) Neighbor-Joining dendrogram based on Nei’s genetic distance. Bootstrap support of nodes is shown. Groups: northern genetic group (Blue); southern genetic group (Red) and c) Scatterplot of the principal components analysis (PCoA) showing the dispersion of samples across the first two principal components

The same two groups identified using the Structure software were also visualized by using the first two PCoA axes as well as the NJ dendrogram. These analyses clearly revealed the formation of two distinct genetic groups within *A. aculeata*, which are suggested to be geographically separated by the Amazon Rainforest (Fig 4b and 4c). Thus, pronounced differentiation was observed between the individuals of *A. aculeata*. Based on the NJ dendrogram, two large groups were assigned based on geographical origin, separating all individuals from the North and Central America in one node (red) from the Brazilian samples (blue). Two main subgroups were evident in the Northern group. One subgroup contained samples from Peritoró (PER) and São Jose dos Patos (SJP) from Maranhão, Brazil. The other subgroup contained the remaining samples. Interestingly, individuals from Tuxtla Chico, Chiapas (TUX) in Mexico formed a separate cluster from the other samples from Mexico and Colombia, Costa Rica, Trinidad and Tobago, and Puerto Rico (Fig 4b).

The second PCoA axis comprised three samples from Cáceres, MT (CAC). These samples formed a subgroup that was very distant from the other samples of *A. aculeata*. However, the Structure and NJ dendrogram data were not able to discriminate these samples, and grouped with individuals from Brazil (Fig 4a and 4c).

The ‘South’ group (Cluster 2 in Fig 4a) contained most of the samples from Brazil. The samples collected in Maranda formed a different cluster from the other samples. However, most clusters reflected a strong relationship with the samples geographic origins, with the exception of samples collected in Belém, PA (BEL), northern Brazil, which were more closely related to samples from Rio de Janeiro and São Paulo located in southeastern Brazil. It is also noteworthy that five samples collected in the Brazilian State of Maranhão (PER and SJP) were more closely related with the ‘North’ group, as evident by the cluster 1 considering the assignment probability of 0.75 in the Structure software analysis (Fig 4a). This result was also corroborated by the NJ and PCoA hierarchical classification (Fig 4b and 4c).

### Genomic diversity of *A. aculeata*

Concerning the genomic diversity within *A. aculeata* species, the greatest diversity was found in Brazil (H_E_ = 0.081) and the lowest diversity in Mexico (H_E_ = 0.005). Likewise, the allelic richness values were similar for all populations in the ‘North’ samples, varying from 1.09 to 1.11. However, the greatest allelic richness for the species was registered in Brazil (Ar = 1.44) (Table 3).

**Table 3.** Genetic diversity parameter estimates for *A*. *aculeata* calculated from 3259 neutral loci of SNPs

| Country | Na | Ne | I | H<sub>O</sub> | H<sub>E</sub> | Ar | PAr | f |
| --- | --- | --- | --- | --- | --- | --- | --- | --- |
| Trinidad &Tobago | 1.001 | 0.983 | 0.013 | 0.008 | 0.008 | 1.09 | 0.01 | 0.038 |
| Colombia | 1.009 | 0.996 | 0.018 | 0.012 | 0.012 | 1.09 | 0.01 | -0.014 |
| Mexico | 0.994 | 0.975 | 0.009 | 0.004 | 0.005 | 1.09 | 0.01 | 0.147 |
| Costa Rica | 0.981 | 0.971 | 0.012 | 0.009 | 0.008 | 1.11 | 0.01 | -0.139 |
| Brazil | 1.441 | 1.124 | 0.135 | 0.043 | 0.081 | 1.44 | 0.33 | 0.377 |
| **Brazilian State** |  |  |  |  |  |  |  |  |
| Pará | 1.090 | 1.044 | 0.059 | 0.046 | 0.039 | 1.11 | 0.01 | -0.168 |
| Mato Grosso | 1.090 | 1.031 | 0.091 | 0.062 | 0.061 | 1.23 | 0.04 | -0.023 |
| Góias | 0.957 | 0.935 | 0.041 | 0.031 | 0.028 | 1.26 | 0.01 | -0.068 |
| Distrito Federal | 0.870 | 0.836 | 0.053 | 0.033 | 0.035 | 1.49 | 0.01 | 0.015 |
| Minas Gerais | 1.190 | 1.094 | 0.089 | 0.043 | 0.058 | 1.09 | 0.01 | 0.198 |
| Rio de Janeiro | 1.024 | 0.987 | 0.037 | 0.024 | 0.024 | 1.14 | 0 | -0.006 |
| São Paulo | 1.200 | 1.089 | 0.091 | 0.044 | 0.059 | 1.10 | 0.01 | 0.179 |
| Maranhão | 1.000 | 0.979 | 0.042 | 0.039 | 0.029 | 1.17 | 0.02 | -0.350 |

Mean of different alleles (Na), effective allele (Ne), Shannon's Index (I), Observed (H<sub>O</sub>) and Expected (H<sub>E</sub>) Heterozygosity, allelic richness (Ar), private alleles richness (PAr) and Fixation index (f)

Due to the vast territory and the greater number of *A. aculeata* samples from Brazil, genetic diversity analyses were conducted considering the Brazilian States as populations. Greater diversity (H_E_) was found in the states of Mato Grosso (MT), São Paulo (SP), and Minas Gerais (MG), with values of 0.061, 0.059, and 0.058, respectively. In terms of allelic richness (Ar), the most accentuated values were located in the central-west region of the country, in Distrito Federal (DF), Goiás (GO), and Mato Grosso (MT), with values of 1.49, 1.26, and 1.23, respectively.

## Discussion

To our knowledge, this is the first study using GBS for identifying genome-wide SNPs and their application for inferring the genetic diversity and population structure in Acrocomia species and within *A. aculeata*. Sampling was broad in terms of the occurrence of Acrocomia species and comprehensively captured the genomic diversity and structure of the species.

### A. aculeata

At the genus level, the distinction of *A. aculeata* as an independent genetic group or taxon was supported through the results obtained with the Bayesian analyses (Fig 2), and by the PCoA and the NJ tree (Figs 3a and 3b). A notable finding was the identification of an accentuated substructure within *A. aculeata*, showing two genetic groups, corresponding to a north– south split in which the samples from Brazil (Northern group, blue cluster in Fig 3) were separated from those of Central and North America (Southern group, red Cluster in Fig 3). This result was evident in the Bayesian analysis performed at the genus level (Fig 2) as well as with only samples of *A. aculeata* (Fig 4a).The substructure identified in *A. aculeata* has not been previously reported and can be attributed to the greater number of samples included in this study, which covered a wide geographic occurrence of the species in the American continent. The presence of two genetic groups may be the result of reproductive isolation due to the Amazon Rainforest acting as a geographical barrier that prevented gene flow between them and with an independent evolution. Another hypothesis is that these two gene pools support the existence of more than one species, as reported in a previous taxonomic classification in Central and North America Countries [57].

Another interesting result observed was that individuals from the population of Maranhão presented as an admixture between the Northern and Southern groups of *A. aculeata* (Fig 4). The origin of the genus Acrocomia is uncertain. However, in the case of *A. aculeata*, based on the dates of archeological records of human use, the most accepted hypothesis suggests that the species originated in northern Brazil (in the region of Santarém, State of Pará) approximately 11,200 MY, and was later dispersed by humans to Central America [58]. According to our results, the admixture observed in the populations of Maranhão (neighboring to Pará State) (Fig 4a) may support this hypothesis, suggesting a common geographical origin of the to two genetic groups in the northeast region of Brazil. In agreement with the *A. aculeata* dispersion routes from South to Central and North America [58], the low values of genetic diversity for the species found in the Northern group may have resulted from a founder effect, since all population of this cluster presented lower values of genetic diversity than those observed in the populations of the southern cluster (Brazil) (Table 3).

Bayesian analysis identified individuals of *A. aculeata* with a degree of genetic admixture with *A. totai* (Cluster 2, in Fig 2) and *A. intumescens* (Cluster 6, in Fig 2), suggesting gene flow between species. As *A. aculeata* is dispersed mainly by cattle [59, 60], the agricultural expansion and livestock may have favored the dispersion of the species to areas where *A. totai* and *A. intumescens* occur, creating opportunities for hybridization due to secondary contact. There have been no reports of interspecific hybridization in the Acrocomia genus. However, a recent study using microsatellite markers also detected connectivity between populations of *A. aculeata* and *A. totai* in Brazil [61].

*A. aculeata* displays the greatest geographical distribution of the genus [3, 4, 14, 15]. As expected for a species with a wide distribution that has adapted to diverse environmental conditions, the genetic diversity of *A. aculeata* was high when compared to other species (Table 3). At the intraspecific level, the highest genetic diversity for the species was found in Brazil, especially in the States of Minas Gerais and São Paulo (Table 3). Although it is not possible to make direct comparisons due to the different types of molecular markers used, previous studies also identified a high genetic diversity for *A. aculeata* in the States of Minas Gerais and São Paulo [23, 28, 29].

An unexpected result was the low genetic diversity of *A. aculeata* in Mexico, where the species is also distributed in an extensive geographical area, from the north to the south of the country (Table 4). These results could reflect the use and exploration of the species in that country and other Central American countries, where adult plants are harvested as the raw material for a fermented drink called “taverna” [62, 63]. This kind of exploration is one of the main factors driving the reduction size or elimination of the natural populations, which affects the reproductive capacity of the species and its natural regeneration [63] and might also been reducing the genetic diversity. *A. aculeata* is strongly associated with humans [58, 59]. Even though it is considered an incipiently domesticated species, it has a wide range of uses in different countries of the Americas [7, 8, 64]. Therefore, patterns of genetic diversity and structure can also be the result of different states of domestication, with different intensities of selection in each region, as also reported for other species, such as beans [65], tomato [66], and cacao [67].

### A. totai

*A. totai* was the second most geographically dispersed species in the genus. It has been documented in eastern Bolivia, Paraguay, Central-west Brazil to northern Argentina [4, 15]. The taxonomic distinction of the species has been demonstrated based on morphological data and geographic distribution [4], leaf anatomy [1], and fruit biometry [68]. However, *A. totai* is commonly regarded as *A. aculeata* due to the pronounced morphological similarity of both species, and because both have fruits with similar biometric and color characteristics [68]. Our results were congruent with the current taxonomic classification of the species. Almost all samples initially considered as *A. totai* (94%) belonged to cluster 2 with a high assignment probability (> 0.75), according to Structure analysis (Fig 2), and corroborated with PCoA and NJ analysis (Figs 3a and 3b). Our results agreed with those of Lima et al. [61], that documented the clear genetic differentiation between *A. aculeata* and *A. totai* (treated as ecotypes) using microsatellite markers. Although not treated as distinct species, but considering the geographical distribution of both, several studies using molecular and morphological markers also reinforced the classification of *A. totai* as a distinct taxon. Lanes et al. (23) used microsatellite markers to demonstrate the marked genetic differences of A. aculeata between individuals from the Pantanal region, State of Mato Grosso do Sul, Brazil, and other regions of the country. Similarly, Silva et al. (27) analyzed the variation in the internal transcribed spacer (ITS) region and identified four haplotypes. Two were shared by genotypes from São Paulo and Minas Gerais, and one was exclusive to genotypes collected in Mato Grosso do Sul. The morphological characteristics of A. aculeata include larger fruits (3.5 and 5.0 cm) and a pulp oil content that can reach approximately 78% (27, 68-70) while the fruits of A. totai are smaller (2.5 and 3.5 cm) with a pulp oil content between 26% and 33% (68, 71, 72.

In Brazil, *A. totai* is considered to be restricted to the State of Mato Grosso do Sul [4, 69, 70]. An interesting finding of our study was that samples from Xambrê, Paraná (XAM) and a sample from Palmas, Tocantins (PAL_182), considered as *A. aculeata* based on Lorenzi et al., [4] taxonomic classification, were attributed to cluster 2 of *A. totai* by the Bayesian analysis (Fig 2), by PCoA, and by NJ (Figs 3a and 3b). Although the occurrence of *A. totai* in these states has not been proven, our results are consistent with the information reported on the Flora do Brazil 2020 website [15], indicating the possible occurrence of *A. totai* in these states.

Although the genetic structure and separation of *A. aculeata* from *A. totai* was evident based on the cluster analyses, the genetic differentiation (F_ST)_ between species was 0.083, which was the lowest value (Table 1). These result was consistent with the value obtained using microsatellite markers (F_CT =_ 0.07) by Lima et al., [61]. The findings may reflect the retention of ancestral polymorphisms, the hybridization or gene flow between species in convergent areas [61] or could be evidence of an ongoing speciation process [23].

Based on the H_E_ and Ar values, *A. totai* was the species with the highest level of genetic diversity (Table 2). Our results are comparable to those found in a recent study using microsatellite markers [61], in which the genetic diversity of *A. totai* was greater than that of *A. aculeata*. Similar, previous studies also identified greater genetic diversity in populations from Mato Grosso do Sul than population from other location of Brazil, although the authors did not consider the populations to be *A. totai* [23, 25]. The high diversity observed in *A. totai* could reflect its geographically widespread occurrence and expansion of genetic diversity promoted by the interspecific hybridization with *A. aculeata*.

The results of cluster analysis and genetic differentiation corroborated the classification of *A. totai* as an independent taxon based on morphological [4], anatomical [1], and molecular markers [61]. This taxonomic separation seems to be more appropriate than that proposed for Henderson et al. [3], which considered all tree-sized Acrocomias as a single taxonomic group called *A. aculeata*.

### A. intumescens

Contrary to the actual taxonomic classification [4–6], our analyses did not show a clear genetic separation of *A. intumescens* (Figs 2, 3a, and 3b). All the samples of *A. intumescens* were assigned to cluster 6, however presented high levels of admixture with *A. aculeata* (cluster 5, Fig 2). *A. intumescens* also showed a moderate genetic differentiation with *A. aculeata* (F_ST_ = 0.128, Table 1), reinforcing the close genetic relationship among both species as described by Vianna et al. [1] based on leaf anatomy. Morphologically, *A. intumescens* is distinguished mainly by the swelling of the stipe [4]. However, botanical characters suggested to delimit Acrocomia species have revealed an overlapping in size of fruits [68] and for oil content in the mesocarp, ranging from 37 to 78% in *A. aculeata* [71, 72] and from 34 to 41% in *A. intumescens* [71, 73].

A phylogenetic study by Meerow et al. [74], estimate the divergence of *A. intumescens* and *A. aculeata* 5 MA ago. The genetic structure we observed may reflect the maintenance of ancestral polymorphism, possibly as a result of the recent divergence of these species with insufficient time for the appearance of reproductive isolation mechanisms, allowing the interspecific hybridization. *A. intumescens* is endemic to northeast Brazil and has a restricted distribution [4, 15]. Species with a restricted geographical distribution tend to have reduced genetic diversity than species with a wide geographical distribution [75, 76]. Consistent with this trend, *A. intumescens* showed lower values of heterozygosity and allelic richness than the wide geographical distribution species *(A. aculeata* and *A. totai*) (Table 2). However, the genetic diversity found in *A. intumesces* was comparable to that observed in other plant species associated with restricted geographic distribution [77–79].

### A. crispa

*A. crispa* is an insular species with a distribution restricted to Cuba. A clear separation and a strong genetic divergence compared to the other species, as evidenced in the cluster analysis (Fig 2, 3a, and 3b) and by the high values of F_ST_ (Table 1). These expectations were understood if considered that the gene flow through pollen or seed dispersal between island populations and continental populations is limited such that a strong genetic structure and a high degree of differentiation between them is expected, as reported for several species [80, 81]. Our results are congruent with those reported for other tree species, which also showed high levels of genetic differentiation between island populations compared to continental populations and lower levels of genetic diversity on the islands than on the continent [82–85]. *A. crispa* displayed low values of genetic diversity (H_E_ = 0.020) compared with other Acrocomia species, although these values are expected for endemic island species. However, interestingly, *A. crispa* presented the greatest allele richness (2.29) and allele richness of private alleles (0.17) (Table 2). Based on chlorosplastic and nuclear genes, the time of divergence estimated for *A. crispa* as 16 Mya, while *A. aculeata* and *A. intumescens* diverged 5 Mya [74]. This more ancient divergence associated with geographic isolation may support the allelic richness and the greater number of private alleles found in *A. crispa*, as well as the strong genetic differentiation of from other Acrocomia species. This hypothesis has also been posited for other endemic species of islands that have congeners on the continent [86, 87].

There is no detailed information about the morphological characteristics of *A. crispa*. However, some morphological differences have been described, such as the presence of swelling in the median region of the stipe as the most discriminating botanical characteristic Bailey [57], the smaller fruits, varies from 1 to 3 cm [3], than that described in *A. totai* (2.5 to 3.5 cm), *A. intumescens* (3.0 to 4.0 cm) and *A. aculeata* (3.5 to 5.0 cm) [68] and also differences in pollen morphology with trichotomocolpated pollen in *A. aculeata* and monocolpous pollen in *A. crispa* (named *Gastrococos crispa* by the authors) [88].

*A. crispa*, previously designated to the genus Gastrococos by Moore [89], was recently allocated to the genus Acrocomia, mainly due to the sequencing of the nuclear *prk* gene [90]. Although most phylogenetic studies that analyzed support the relationship between *A. aculeata* and *A. crispa* as sisters in a single monophyletic group [90–95], other phylogenetic [74] and cladistic studies [96] shown that they are sister species in paraphyletic groups. However, these phylogenetic studies were conducted at higher taxonomic levels (families, subfamilies, and tribes), with the inclusion of few species of Acrocomia. Therefore, they have limited ability to accurately reveal phylogenetic relationships of Acrocomia species.

The morphological characteristics of the species, the divergence time and our results of genetic differentiation, diversity, and structure may collectively support an independent taxonomic status of *A. crispa* within the genus Acrocomia. Therefore, we suggest a revision of the taxonomy for the species.

### A. media

In contrast to the evidence of genetic divergence for *A. crispa*, the recognition of *A. media* as an independent taxonomic unit was not supported by our study. As *A. media* is also an island species, it would be expected to have a strong genetic structure when compared to other Acrocomia species with a continental distribution. Contrary to this assumption, all samples considered as *A. media* were assigned to the northern group of *A. aculeata*, as evidenced by three cluster analyses (Figs 2, 3a, and 3b). In addition, the F_ST_ values (Table 1) also indicated low genetic differentiation of *A. media* compared to *A. aculeata*.

The patterns of genetic diversity observed in *A. media* were the lowest compared to other species (H_E_= 0.005 and Ar = 1.08), but were consistent with several studies of population genetics in plants, which predicted that island populations have reduced levels of genetic diversity compared to continental populations [80, 97]. The low genetic diversity observed in *A. media* can be attributed to the founder effect associated with the establishment of populations with only a few individuals [97, 98] or to genetic drift due to stochastic events inherent in the islands and/or fragmentation during its formation [99]

*A. media* was first described in Puerto Rico by Cook [100]. The author adopted the shortest trunk and the smallest diameter of the stipe as the differentiating characteristics of *A. media* from *A. aculeata*. However, *A. media* was considered synonymous with *A. aculeata* for a long time due to the absence of consistent botanical characteristics for differentiation. In 2013, The Plant List recognized *A. media* as a distinct species based on the floristic palm inventory of Proctor [101]. However, the same author mentioned that the existing information about *A. media* was very old and based on few individuals, suggesting an increase in the number of evaluated individuals to guarantee a more consistent morphological description of the species. The only phylogenetic study performed with *A. media* included an individual from Puerto Rico, and a sample of *A. aculeata* from Brazil revealed that both species were closely related [90].

Based on the lack of genetic differentiation of *A. media*, low genetic diversity in the species, and low pairwise F_ST_ value between *A. media* and *A. aculeata*, we hypothesize that *A. media* is synonymous with *A. aculeata*. Thus, a recent introduction in Puerto Rico was not sufficient to characterize the reproductive isolation needed for the differentiation of *A. aculeata*.

### A. hassleri and A. glauscescens

The genomic data of our study did not allow the assignment of distinct taxonomic units to the species *A. hassleri* and *A. glauscescens*. Based on morphological characters, the species are clearly differentiated from the others by their small size. However, based on the results obtained from the cluster analysis, they were assigned to cluster 2, being closely related to *A. totai* (Fig. 2, 3a, and 3b). However, this result should be considered with caution, as we only used one sample of each species in the analyses, which could limit the comparison of genetic estimates and decrease the probability of detecting genetic structure, as evidenced in similar studies with a low number of samples [102, 103]. Further studies with a greater number of accessions are needed to increase the species representation, and to establish reliable genetic relationships between *A. hassleri* and *A. glauscescens* and other Acrocomia species.

## Conclusions

Our study is the first to offer evidence of the efficiency of NGS through the application of the GBS protocol in Acrocomia. The data may constitute a reference for the application of this protocol in the genus. Even without a reference genome, we successfully identified a large number of SNPs for several species, revealing potentially valuable markers for future studies in the genus Acrocomia. The SNPs yielded unprecedented results of the genetic relationships between Acrocomia species as well as at the population level for *A. aculeata*. In general, our results were partially congruent with the taxonomy of the genus, supporting the current separation of some species. The genomic structure revealed the formation of well-defined genetic groups and confirmed the distinction of *A. aculeata, A. totai, A. intumescens*, and *A. crispa*, with the latter showing a strong genetic differentiation as well as the absence of genetic distinction of *A. media*. We recommend a review of the current taxonomic classification of *A. crispa* and *A. media*. In addition, SNPs also allowed the identification of gene flow patterns and/or hybridization between species.

In the case of *A. aculeata*, the data provide an overview of the genomic diversity and structure from sampling over a wide area of occurrence. The genomic data showed the existence of two large gene pools in the species at the continental level (north and south), with greater genomic diversity in the latter populations. The results from this study will serve as a reference for current and future studies on genetic diversity, taxonomy, evolution, ecology, and phylogeny of the genus Acrocomia, and will support genetic breeding, conservation, and management activities for *A. aculeata*.

## Supporting information

**S1 Table. Geographical location and origin of the Acrocomia species samples**.

**S2 Table. SNP loci outliers (putatively under selection) for Acrocomia species and within *A*. *aculeata* identified by PCAdapt, Fsthet and LEA packages**.

**S1 Fig. Delta (Δ) K values for different numbers of populations assumed (K) in the STRUCTURE analysis, estimated based on Evanno method for all Acrocomia species**.

**S2 Fig. Delta (Δ) K values for different numbers of populations assumed (K) in the STRUCTURE analysis, estimated based on Evanno method for *A*. *aculeata***.

**S1 File. SNP genotype information in variant calling format (vcf) for 172 samples of Acrocomia species**.

